# miR-92a-3p controls cell cycle progression in zebrafish

**DOI:** 10.1101/680991

**Authors:** Christopher E. Presslauer, Teshome T. Bizuayehu, Jorge M.O. Fernandes, Igor S. Babiak

## Abstract

Biological functions of micro RNAs (miRNAs) in the early stages of vertebrate development remain largely unknown. In zebrafish, miRNA miR-92a-3p is abundant in the germ cells throughout gonadal development, as well as in ovulated oocytes. Previously, we demonstrated that inhibition of miR-92a-3p in mature ovaries resulted in developmental arrest at the 1-cell stage upon fertilization of the affected oocytes. This suggested functions of miR-92a-3p in early development. In the present study, we identified *wee2*, an oocyte-specific protein tyrosine kinase, as a target of maternal miR-92a-3p during the early stages of zebrafish embryogenesis. Spatiotemporal co-presence of both miR-92a-3p and *wee2* during early embryo development was confirmed by absolute quantification and *in situ* hybridization. Targeted knockdown of miR-92a-3p in embryos resulted in retarded embryonic development over the first 24 hours. Target validation assays demonstrated that miR-92a-3p interacted with the predicted *wee2* 3’UTR binding site, which was strongly suppressed by endogenous miR-92a-3p. Our results suggest that miR-92a-3p regulates the abundance of *wee2*, a cyclin-dependent kinase 1 inhibitor, thus having important role in regulation of the cell cycle during cleavage stages in zebrafish.

**Summary statement:** In zebrafish, maternal miR-92a-3p was demonstrated to suppress translation of *wee2*, a cyclin-dependent kinase 1 inhibitor which regulates cell cycle progression during the early stages of embryogenesis.

## Introduction

MicroRNAs (miRNAs) are a family of small non-coding RNAs which function in post-transcriptional repression of protein-coding genes (Bartel 2009). A miRNA along with Argonaute (Ago) protein is an essential component of the miRNA-induced silencing complex (miRISC), where it acts as the target recognition component (Ha and Kim 2014; Li and Rana 2014). The target recognition is primarily based on base-pair complementarity between nucleotides 2-8 of the miRNA (the seed region) and the 3’ untranslated region of targeted mRNA, resulting in translational repression, mRNA de-adenylation, or mRNA decay (Huntzinger and Izaurralde 2011; Ha and Kim 2014).

The biological functions of miRNAs during teleost gametogenesis and further early embryonic development are poorly understood (Bizuayehu and Babiak 2014). miRNAs are abundant in ovulated oocytes of teleosts (Ma *et al.* 2012; Ma *et al.* 2015; Presslauer *et al.* 2017), and newly formed embryos are transcriptionally quiescent; their development is driven by maternally provided proteins and RNAs (Tadros and Lipshitz 2009; Langley *et al.* 2014; Lee *et al.* 2014b). While maternal miRNAs are known to function in model species such *Drosophila melanogaster* (Marco 2015), *Caenorhabditis elegans* (McJunkin and Ambros 2017), or mouse (Tang *et al.* 2007a), to date only miR-202-5p has been confirmed as a maternal miRNA in teleost embryos, but its maternal function remains elusive (Zhang *et al.* 2017).

Currently, miRNA target prediction is based primarily on computational algorithms, which perform alignments between miRNA sequences and regulatory regions of mRNAs (John *et al.* 2004; Krek *et al.* 2005; Friedman *et al.* 2009). However, these methods often generate many false positive predictions (Ekimler and Sahin 2014) and often do not take into account whether a miRNA and its predicted target are co-expressed (Ritchie *et al.* 2009). It is therefore important that miRNA/mRNA target interactions are verified experimentally.

Recently, we profiled miRNA transcriptome throughout the gonadal development of zebrafish (Presslauer *et al.* 2017). miRNA miR-92a-3p was consistently among the most abundant miRNAs in developing and mature gonads of both sexes. Notably, in ovulated eggs, size variants of miR-92a-3p were dominant, particularly those with one to three additional templated adenine residues (Presslauer *et al.* 2017). Adenylation of maternal miRNAs in oocytes has been reported in *D. melanogaster*, sea urchin (*Stronglyocentrotus purpuratus*), and mouse (Lee *et al.* 2014a), suggesting miR-92a-3p may also be stored in zebrafish oocytes with a maternal function in embryos. Previously, we demonstrated that targeted knockdown of miR-92a-3p in mature zebrafish ovaries resulted in a significant reduction of miR-92a-3p in 1-cell embryos, accompanied by a significant increase in the proportion of embryos with arrested development at the 1-cell stage (Presslauer *et al.* 2016).

In the present study, we examine the spatial and temporal expression patterns of miR-92a-3p in zebrafish gonads and early embryonic development. We identify *wee1 homolog 2* (*wee2*) as a target for maternal miR-92a-3p, and demonstrate miRNA/mRNA co-expression and interactions.

## Results

### miR-92a-3p target prediction

Twenty-five protein families within the zebrafish cell cycle pathway were identified (Supplementary File S1). Noticeably, the predicted targets were within components involved in DNA biosynthesis, such as the mini-chromosome-maintenance complex and origin recognition complex. In addition, a number of predicted targets are involved in the cyclin dependent kinase (Cdk1) pathway at the G2 / M checkpoint of the cell cycle (Fig. 1). From these predicted Cdk1 pathway targets, *wee1 homolog 2* (*wee2*) was selected for target validation assays, based on its confirmed maternal abundance in zebrafish (Aanes *et al.* 2011).

**Figure 1.**
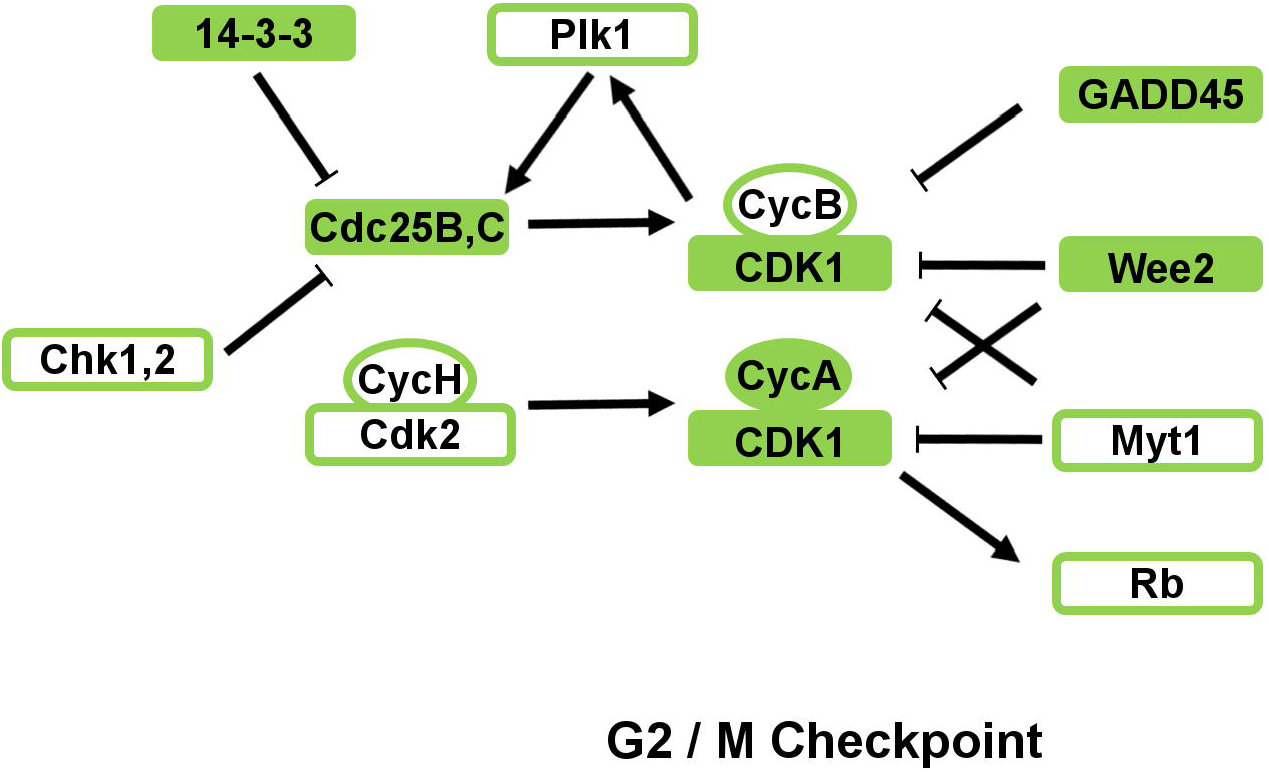
Predicted miR-92a-3p targets within the Cdk1 regulatory pathway. Figure was modified from the KEGG pathway for the zebrafish cell cycle (Kanehisa and Goto 2000). Filled boxes indicate proteins with transcripts predicted as targets for miR-92a-3p, whereas empty boxes indicate non-targets.

### Spatiotemporal expression in gonads and embryos

The germ cells in 10-week-old zebrafish testis predominantly consisted of spermatogonia, spermatocytes, and spermatids; whereas in the ovaries, they consisted of stage Ib and II oocytes, with some stage Ia oocytes located at the periphery. Hybridization with a miR-92a-3p-specific probe identified transcripts in cells of both germline and somatic origin (Fig. 2), with staining particularly strong in primary and vitellogenic oocytes.

**Figure 2.**
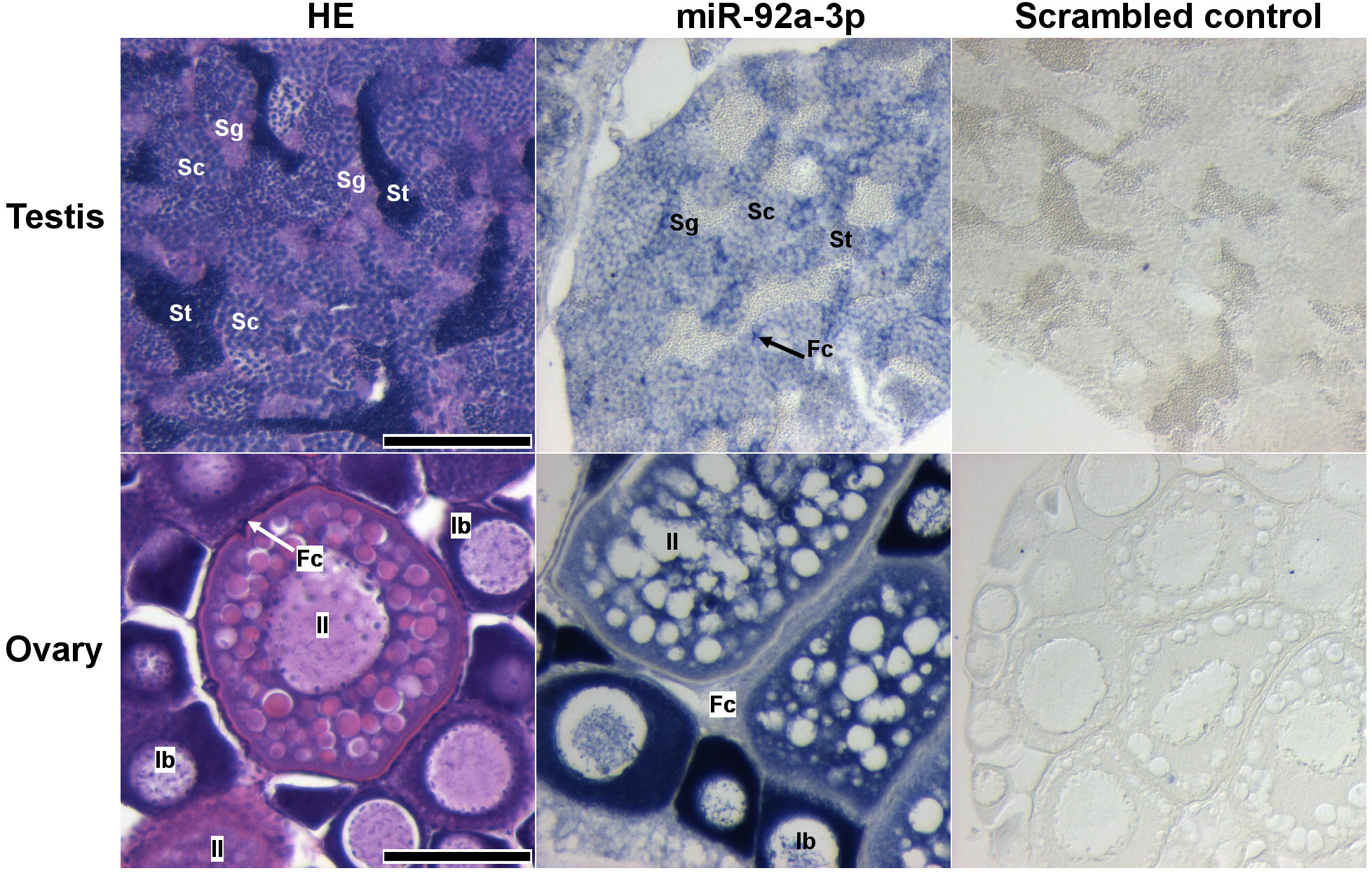
*In situ* hybridization of miR-92a-3p in 10-week-old zebrafish testes and ovaries compared with hematoxylin and eosin (HE) stained sections and sections hybridized with a scrambled negative control. Sg = spermatogonia, Sc = spermatocytes, St = spermatids, Fc = follicle cells, Ia = primary growth pre-follicle stage, Ib = primary growth follicle stage, II = cortical alveolus stage. Scalebars represent 100 μm.

Both miR-92a-3p and *wee2* transcripts were detected in zebrafish embryos. Mature miR-92a-3p transcripts were strongly expressed in zebrafish embryos at all stages examined (Fig. 3 A). At the 1-cell and 256-cell stages, the transcripts were detected ubiquitously within the cells. At the 25 somite stage, miR-92a-3p transcripts were predominantly localized to the eyes and brain, but the signal was detected throughout the body. Expression within the somites was comparatively weaker than in the brain but consistent, while staining was noticeably reduced in the notochord. By comparison, transcripts of *wee2* appeared to be strongly expressed at the 1-cell stage. But the signal appeared weaker at the 256-cell stage (Fig. 3 B). At the 25 somite stage, a faint signal was only detected in the brain and ventral portion of the somites. By comparison, no signal was detected in 1-cell embryos hybridized with negative control probes.

**Figure 3.**
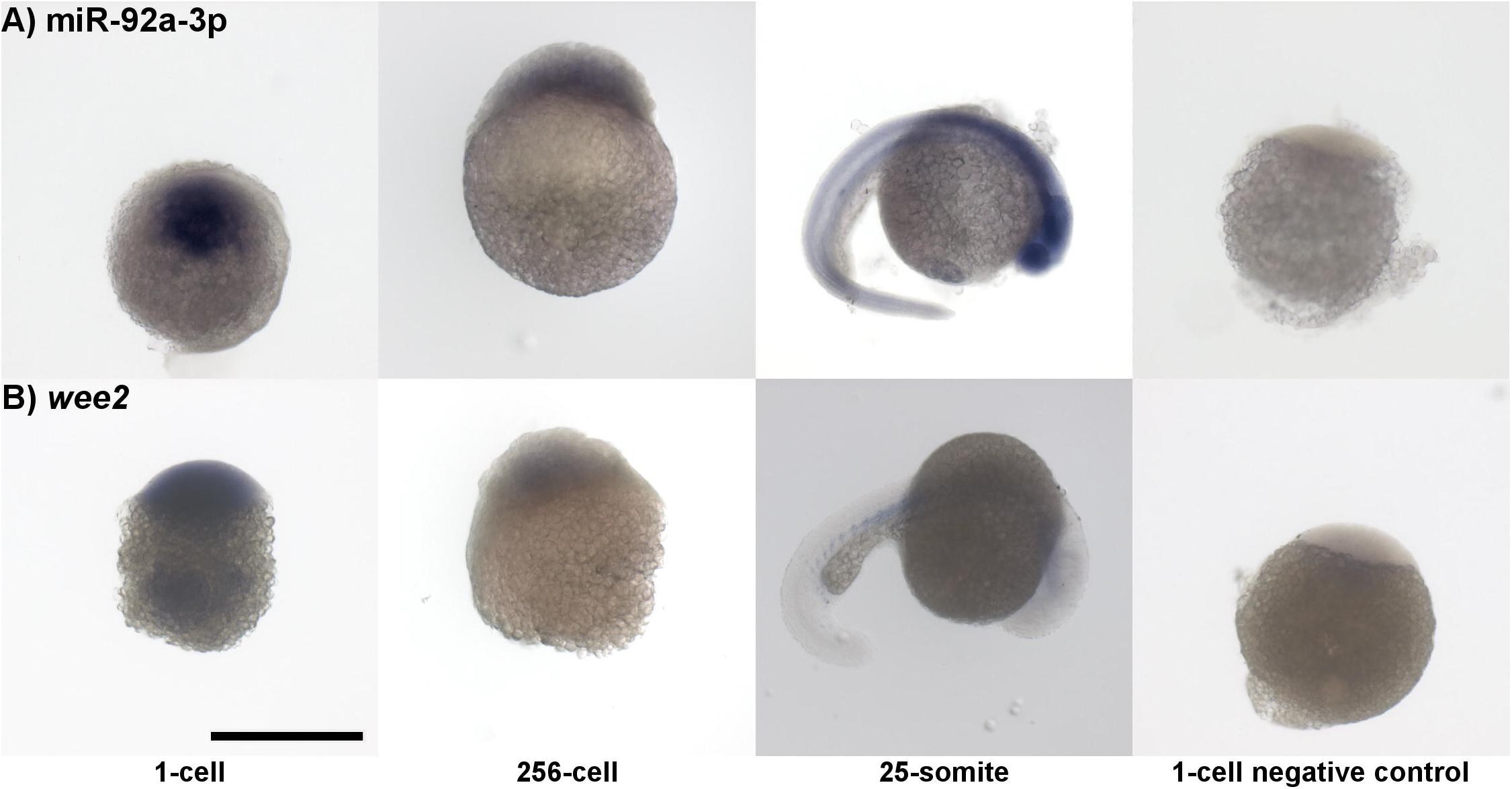
Whole mount *in situ* hybridization of miR-92a-3p (A) and *wee2* (B) at the 1-cell, 256-cell, and 25-somite stages. The negative control for miR-92a-3p is a LIMA™ Scramble-miR negative control probe, while for *wee2* the antisense sequence of the gene specific probe was used. The scalebar represents 500 μm.

Transcripts of miR-92a-3p and *wee2* were highly abundant in early stage embryos, with stable levels observed between activated embryos and the oblong stages of development (Fig. 4). At the 25 somite stage, *wee2* transcripts were barely detectable; in contrast, miR-92a-3p had a significant increase in transcript abundance. Complete droplet digital PCR results with reference genes are available in Supplementary File S2.

**Figure 4.**
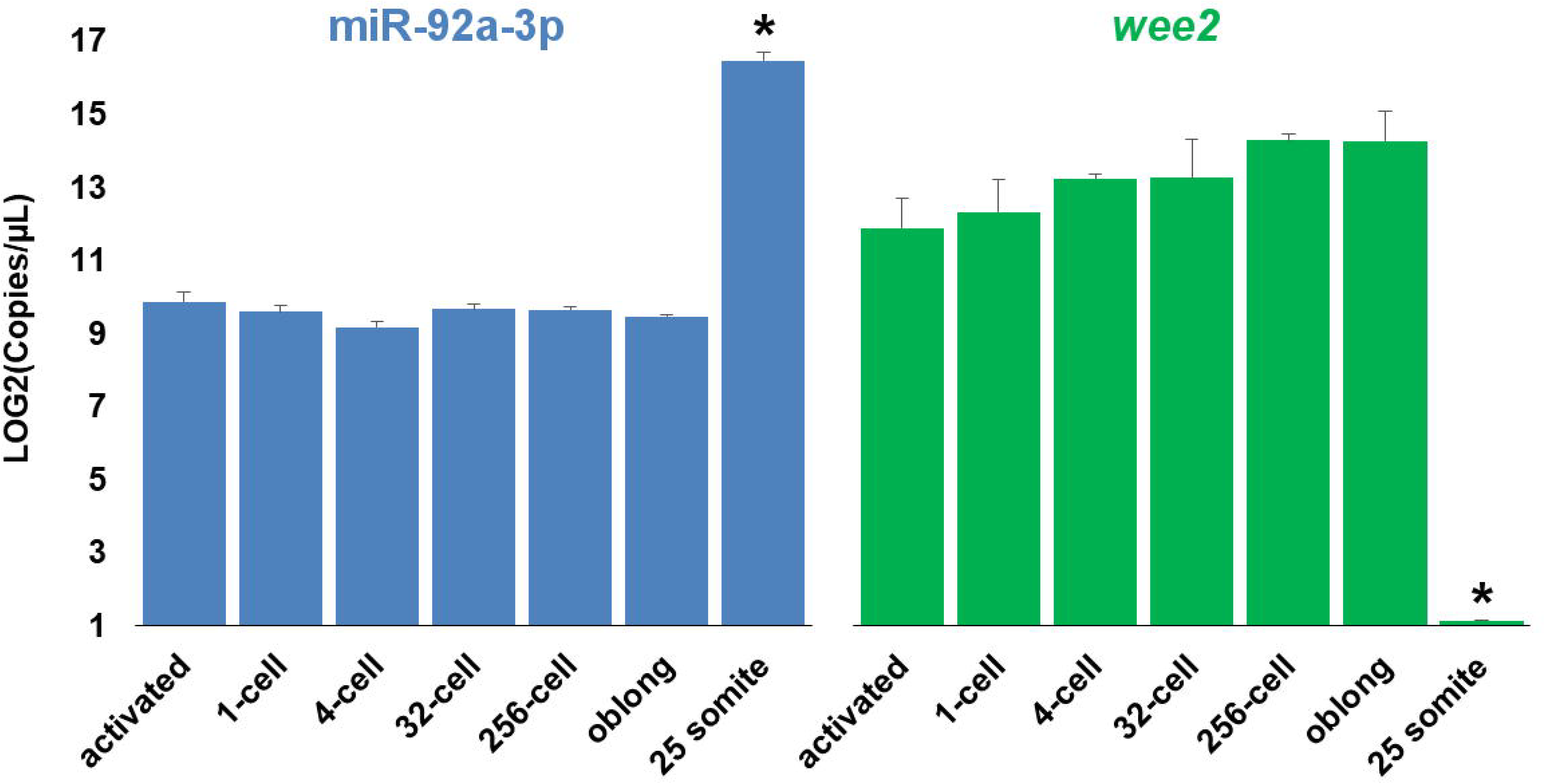
Droplet digital PCR of miR-92a-3p and *wee2* RNAs in zebrafish embryos. Values represent the log2-transformed number of transcript copies per μL cDNA input. Error bars are the standard deviation of the log transformed values. Within a transcript, asterisk indicates a significant difference (*p* ≤ 0.05) from all other developmental stages.

### *gfp-wee2* reporter expression in embryos

Injections of 200 pg or 800 pg *gfp-wee2* resulted in no visible GFP signal (Fig. 5 A), similar to non-injected control embryos (Fig. 5 B). Injections of 1600 pg mRNA produced slightly elevated GFP signal within the yolk, while the GFP signal within the body was barely detectable. An increased concentration of 3200 pg *gfp-wee2* resulted in consistent and strong expression of GFP in the embryo (Fig. 5 A). Notably, throughout the experiment, embryos with severe deformities produced intense GFP signal from reduced mRNA concentrations (Fig. 5 C). By comparison, embryos which received injections of chimeric mRNA containing the antisense sequence for the zebrafish *wee2* 3’UTR had a strong and ubiquitous GFP signal, which was easily visible from concentrations of 200 pg mRNA onwards, and highly intense at 1000 pg (Fig. 5 D).

**Figure 5.**
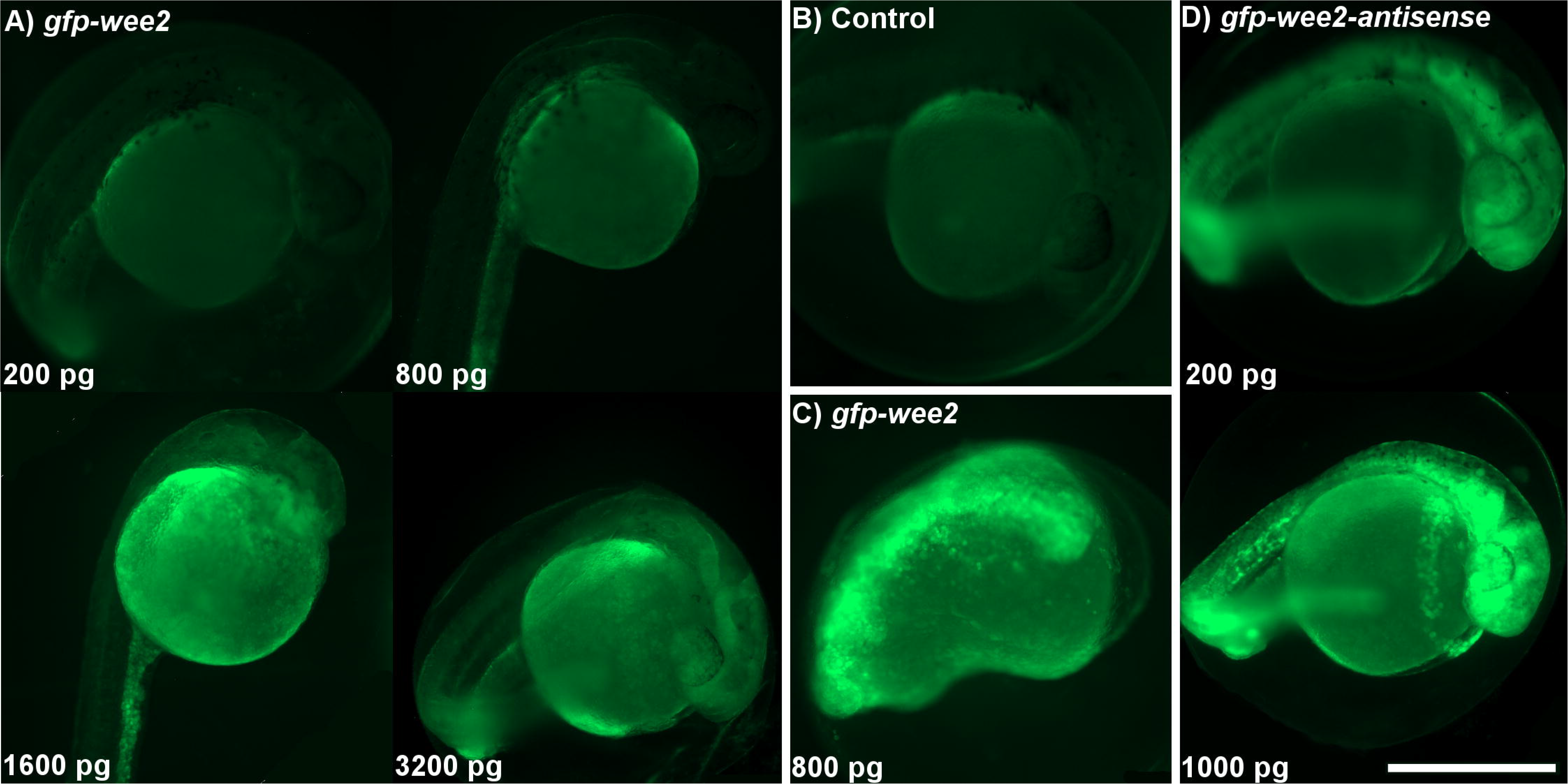
GFP-3’UTR chimeric RNA injections into zebrafish embryos. **A)** Representative images of zebrafish embryos injected with 200, 800, 1600, or 3200 pg of *gfp-wee2* mRNA. GFP signal was detected within the trunks of embryos injected with 1600 or 3200 pg *gfp-wee2* mRNA, but not in embryos injected with lower concentrations. B) A non-injected control embryo. C) A representative image for *gfp-Wee2*-injected embryos showing developmental malformations. D) Representative images of zebrafish embryos injected with 200 or 1000 pg *gfp-wee 2-anti sense* mRNA. All observations were made at 1 dpf. The scalebar represents 500 μm.

### miR-92a-3p target site blocker and *gfp-wee2* co-injections

Co-injection with *gfp-wee2* and either a scrambled control or miR-92a-3p target site blocker (TSB) resulted in varying proportion of GFP expressing embryos and the relative intensity of the GFP signal produced (Fig. 6). Across the three trials, a total of 36 and 52 embryos were injected with *gfp-wee2* and either the control TSB, or TSB specific to the miR-92a-3p predicted target site, respectively. In total, only six out of 36 embryos (6/36) receiving the control TSB had detectable GFP signal, compared to 35/52 receiving the miR-92a-3p TSB (Fig. 6 A). In addition, the mean GFP signal intensity in embryos receiving the control TSB was significantly lower than in those receiving the miR-92a-3p TSB (Fig. 6 B; representative embryos in Fig. 6 C).

**Figure 6.**
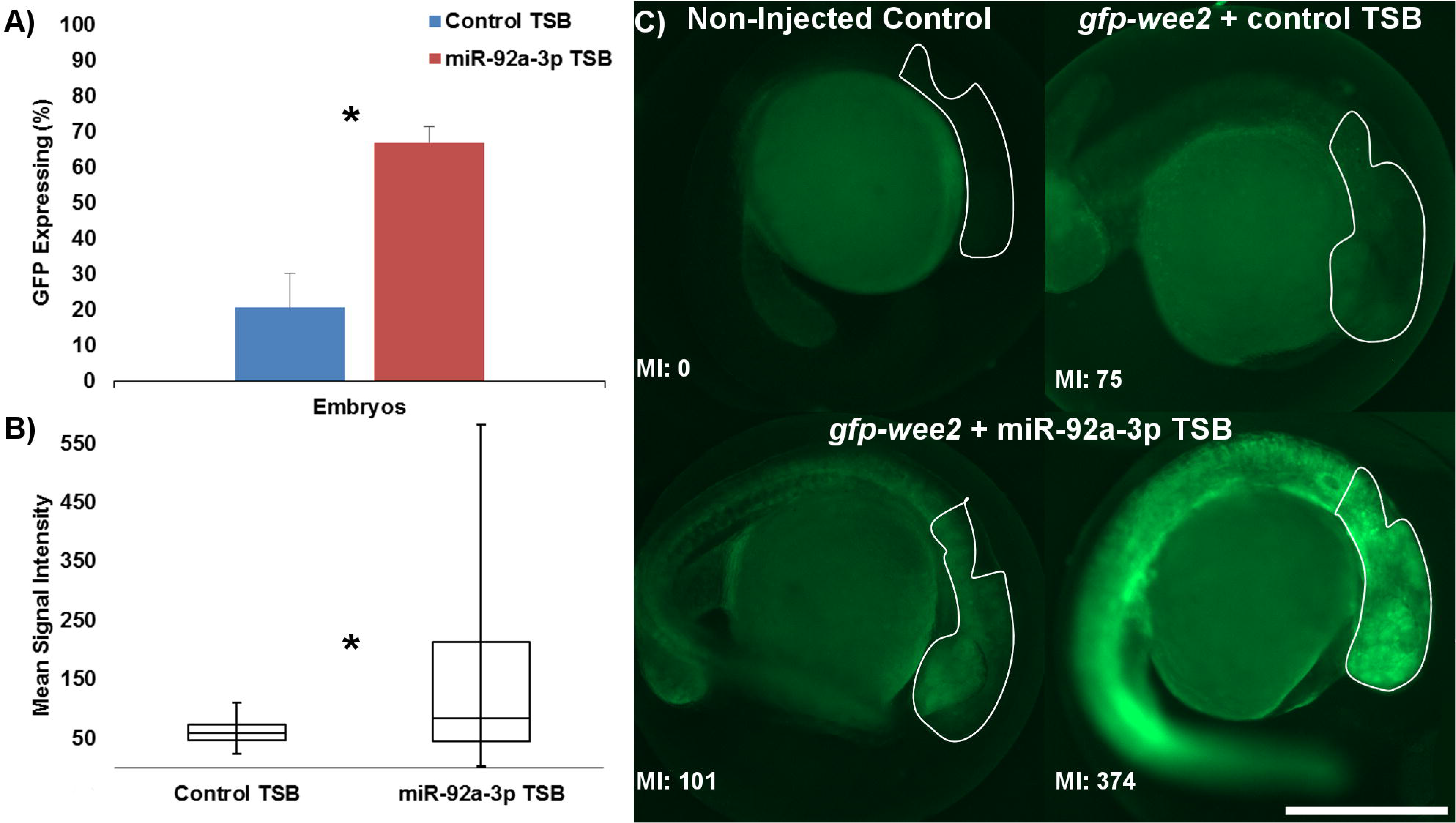
Zebrafish miR-92a-3p target protector assays. Microinjections of *gfp-wee2* chimeric RNA with a target site blocker (TSB) specific for the miR-92a-3p predicted binding site were made to determine miRNA/mRNA interaction. A) The relative proportion of embryos with detectable GFP signal between the control TSB and miR-92a-3p TSB treatments. B) Box plots display the distribution of GFP mean intensity values from all embryos in the two treatment groups. Asterisks mark significant differences at *p* ≤ 0.05. C) Representative embryos from the non-injected control, *gfp-wee2* and control TSB, and *gfp-wee2* with miR-92a-3p TSB treatments. White contours identify the area of the zebrafish head in which the GFP mean intensity (MI) value was measured (values displayed bottom left). The scalebar represents 500 μm.

### miR-92a-3p mimic and *gfp-wee2* co-injections

Co-injections of *gfp-wee2* reporter at variable doses (800 pg to 3200 pg) with miR-92a-3p mimic at a constant dose (5.0 μM) demonstrated that the effect of the mimic was overcome only with the high doses of the reporter (Fig. 7 A). Only 2/9 embryos injected with 800 pg *gfp-wee2* mRNA showed the GFP signal, while no GFP signal was observed in embryos co-injected with the miR-92a-3p mimic (0/5). For the dosage 1600 pg, the number of GFP expressing embryos was 19/25 and 1/12, respectively, and for the dosage 3200 pg it was 9/10 and 7/8, respectively, for injections with the reporter and co-injections with the reporter and mimic. While injections of 3200 pg of the reporter were sufficient to overcome the mimic and produce a GFP signal, the GFP intensity was reduced, although not significantly due to the high interspecific variability (Fig. 7 B; representative GFP expressing embryos in Fig. 7 C).

**Figure 7.**
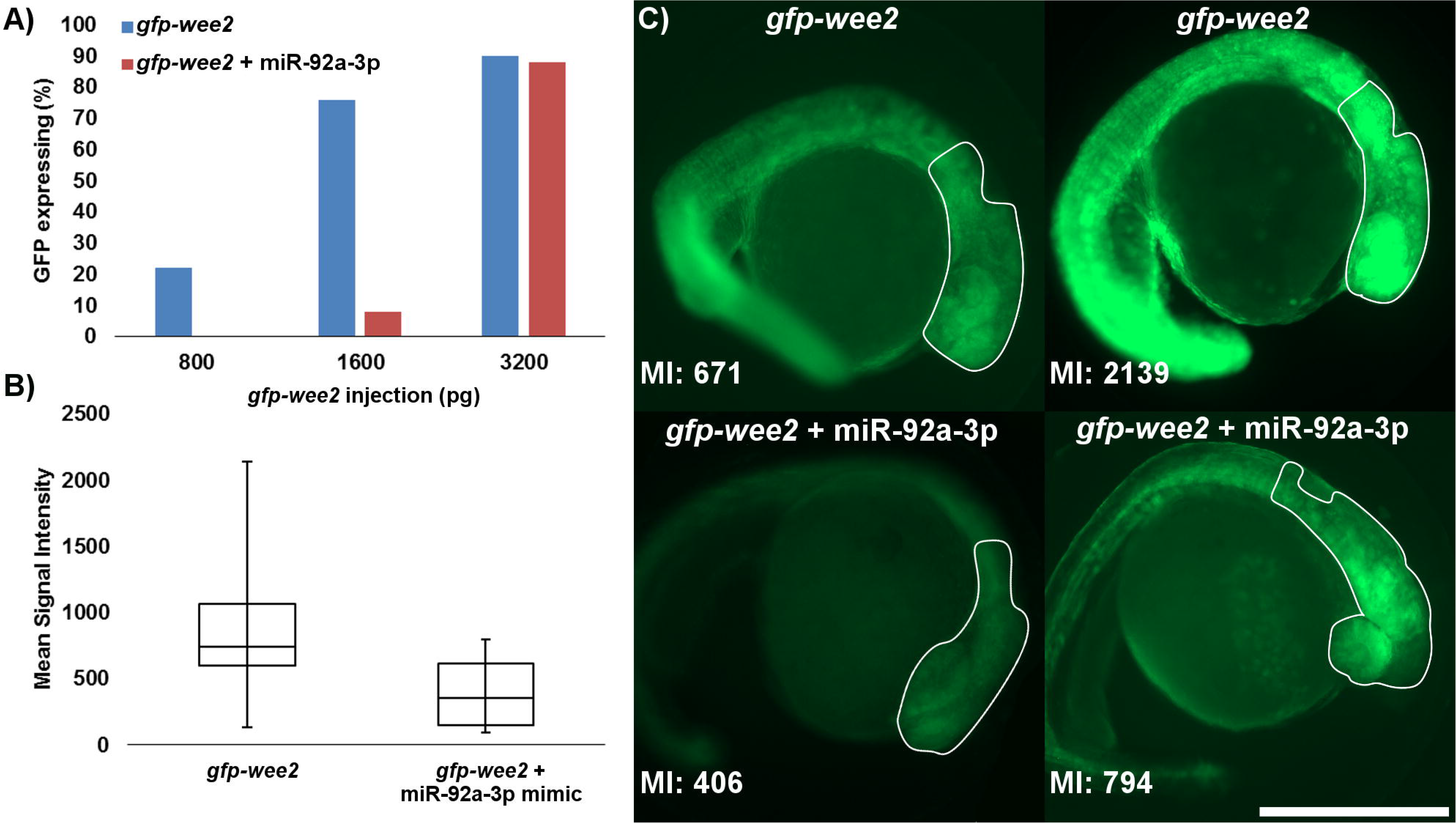
Zebrafish miR-92a-3p target validation assays. Microinjections of *gfp-wee2* chimeric RNA with synthetic miR-92a-3p mimic to assess miRNA/mRNA interactions. A) The proportion of embryos with detectable GFP signal after injections with various doses of *gfp-wee2* mRNA, and those co-injected with miR-92a-3p mimic (5.0 μM). B) Box plots display the distribution of GFP mean intensity values between embryos receiving 3200 pg of *gfp-wee2* mRNA, and those also receiving the miR-92a-3p mimic co-injection. Horizontal lines represent the quartiles, with the center horizontal line representing the mean value. C) Representative embryos injected with *gfp-wee2* mRNA (3200 pg) alone, or co-injected with both *gfp-wee2* and miR-92a-3p mimic. White contours identify the area of the zebrafish head in which the GFP mean intensity (MI) value was measured (values displayed bottom left). The scalebar represents 500 μm.

### miR-92a-3p knockdown and rescue

The majority of miR-92a-3p knockdown embryos (23/39) showed delayed development, defined as inability to reach at least the 20-somite stage by 1 day post-fertilization (dpf); one of these embryos reached only the 1-somite stage. By comparison, all the wildtype embryos reached the 25-somite stage by 1 dpf. A low proportion of morphants were in both rescue and mismatch MO control treatments (4/47 and 2/40, respectively), and no morphants were observed in the miR-92a-3p mimic group (0/39; representative phenotypes given in Fig. 8 A).

**Figure 8.**
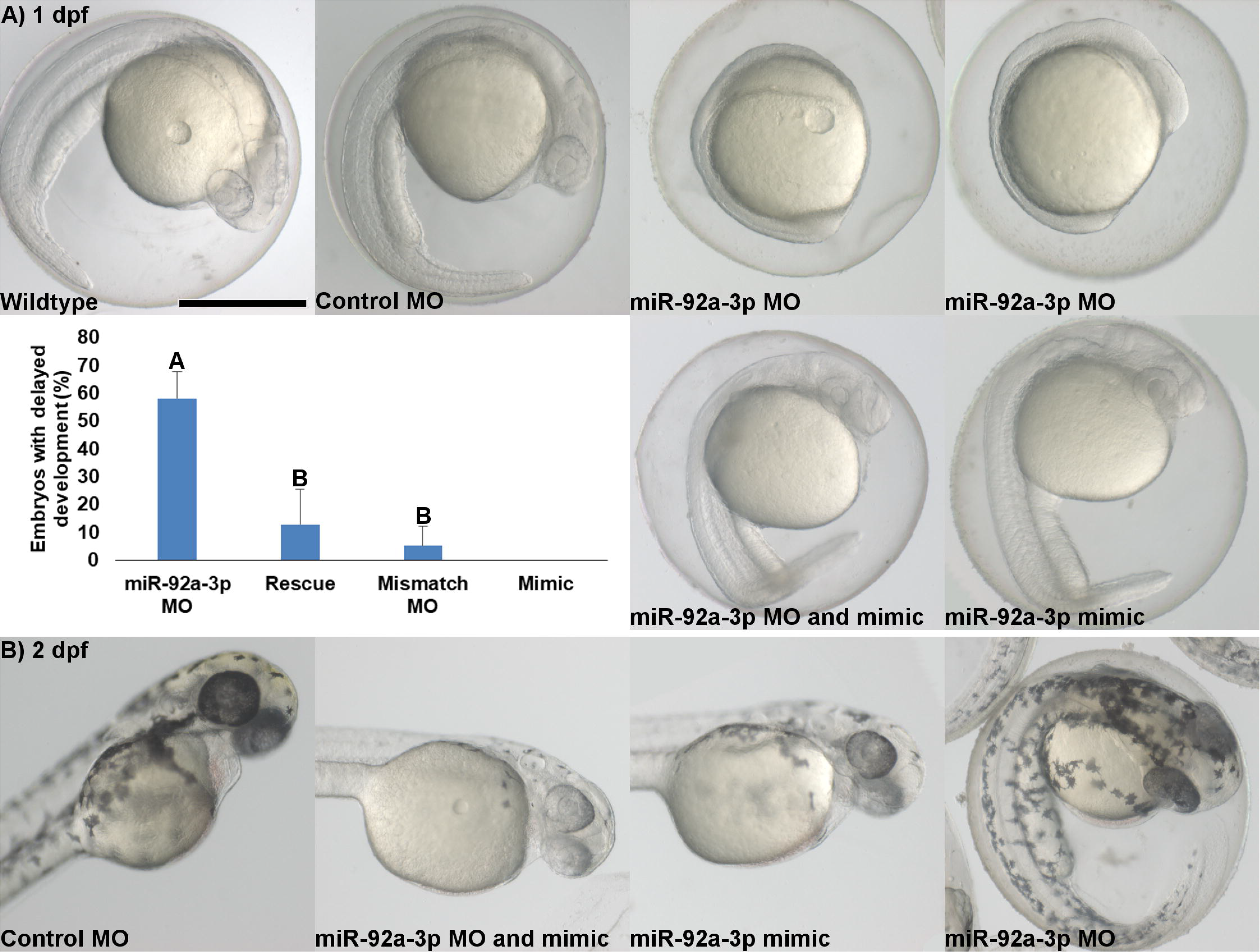
miR-92a-3p knockdown and rescue in zebrafish embryos. A) Representative images display typical development of zebrafish embryos at 1 day post fertilization after the treatments. Wildtype (non-injected control) embryos are at the 25-somite stage of development, similar to typical embryos which received a scrambled MO, a miR-92a-3p MO and miR-92a-3p mimic co-injection, and a miR-92a-3p mimic only. However, most of miR-92a-3p MO embryos showed retarded development. The chart displays proportion of embryos in each treatment group with the delayed development, as defined by not having reached the 20-somite stage, observed at 1 day post fertilization. Different letters identify groups significantly different from each other between the treatment groups (*p* ≤ 0.05). B) Follow up observations at 2 days post fertilization. Embryos from mimic and MO + mimic treatments show malpigmentation and slight developmental delay, as compared to control and MO embryos.

At 2 dpf, embryos from the mismatch MO control group reached the long pec stage of development, while rescue and miR-92a-3p mimic embryos generally displayed slightly delayed development and appeared closer to the high pec stage of development. The rescue and miR-92a-3p mimic treatments often displayed reduced pigmentation when compared to the MO only treated embryos. Interestingly, the miR-92a-3p MO-injected embryos, which suffered severe retardation in development at 1 dpf, had recovered at 2 dpf. They were at the high pec stage of development and exhibited a normal pigmentation pattern, similar to controls (Fig. 8 B).

## Discussion

### miR-92a-3p and its predicted target *wee2* are abundant maternal factors in zebrafish

Previously we have found miR-92a-3p among the most abundant miRNAs throughout the gonadal development in zebrafish and massively represented in mature oocytes (Presslauer *et al.* 2017). Also, Vaz *et al.* (2015) have found this miRNA abundant in mature gonads of zebrafish. The present study demonstrates that the gonadal localization of miR-92a-3p transcripts is in germ cells (Fig. 2) and embryonic localization is within the cell prior to the first cleavage (Fig. 3), providing further evidence of maternal inheritance. Both WISH and ddPCR methods detected high abundance of miR-92a-3p throughout embryogenesis (Figs. 3, 4). The significant increase in miR-92a-3p abundance between the oblong and 25 somite stages (Fig. 4) indicates zygotic transcription as its source in later developmental stages. This pattern differs from the previous reports on zebrafish where miR-92a-3p was suggested to be first expressed during the blastula stage (Li *et al.* 2011), or where miR-92a-3p gave relatively weak signal at 1 dpf, before it greatly increased in expression at 2 and 3 dpf (Ning *et al.* 2013). However, evidence obtained in zebrafish (Presslauer *et al.* 2016; Presslauer *et al.* 2017) and rainbow trout (*Oncorhynchus mykiss*), another teleost fish (Ma *et al.* 2012), rather clearly suggests that miR-92a-3p is a maternally-inherited miRNA with functions in the zebrafish embryo prior to the maternal-zygotic transition, and its expression in the later development is of zygotic transcription origin.

miR-92a-3p is a member of the conserved miR-17-92 cluster, which functions in regulating mammalian cell cycle, proliferation, and apoptosis (Manni *et al.* 2009; Mogilyansky and Rigoutsos 2013; Zhou *et al.* 2015). In human cell lines, miR-92a-3p was determined to function in promoting the cell cycle transition from the G1 to S phase (Zhou *et al.* 2015). In zebrafish, miR-92a-3p is most strongly expressed in proliferative organs such as the brain, suggesting its role in regulating cell cycle and proliferation is a conserved function (Ason *et al.* 2006; Ramachandra *et al.* 2008). Within the zebrafish cell cycle pathway, we identified a number of predicted targets for miR-92a-3p, including *wee2* (Fig. 1, Supplementary File S1). Wee2, a tyrosine kinase, was identified in *Xenopus* as a Cdk1 inhibitor capable of stopping cell cycle progression (Leise and Mueller 2002; Leise and Mueller 2004). Its role as an oocyte-specific key regulator of meiosis in prophase I and metaphase II through inhibition of CDK1/CDC2 is conserved in mammals, and its deficit leads to fertilization failure in human (Hanna *et al.* 2010; Sang *et al.* 2018). The metaphase II arrest is common in vertebrates, including mammals and fish, and its mechanism, termed Cytostatic Factor, is conserved (Madgwick and Jones 2007). However, functions of Wee kinases exert beyond oocyte-specific processes. As demonstrated in *Xenopus*, the Wee family kinases *wee1, wee2*, and *myt1* have unique spatiotemporal expression patterns during embryogenesis (Leise and Mueller 2002). Notably, while *wee1* and *myt1* are expressed maternally, *Xenopus wee2* is only expressed after the maternal to zygotic transition, where it is predominantly found in locations lacking proliferating cells. However, injections of either *wee2* mRNA or protein into 2-cell embryos resulted in delay or arrest of cleavage (Leise and Mueller 2002).

In zebrafish, *wee2* transcripts are among the most abundant maternal mRNAs during the early stages of embryonic development (Aanes *et al.* 2011). Our ISH results were consistent with the study by Aanes et al. (2011), in that *wee2* was strongly expressed during the cleavage stage, before declining in the blastula stage, and was barely detectable during somitogenesis (Fig. 3 B). The ddPCR results differed in that expression remained high at the oblong stage before it ceased by the 25-somite stage (Fig. 4). Both methods indicate that *wee2* is a maternally-provided transcript in zebrafish early embryos. In addition, the co-expression of *wee2* and miR-92a-3p suggest there is a strong potential for interaction between this miRNA and its potential target.

### *wee2* is inhibited by endogenous factors

*wee2* is not among the predicted targets of the miR-430 family (Vejnar and Zdobnov 2012; Agarwal *et al.* 2015), a major clearance factor of maternal mRNA after maternal-to-zygotic transition (Giraldez *et al.* 2006). The decline of *wee2* transcript level after the maternal-to-zygotic transition suggests that other factors can be involved in its degradation.

Injections of a *gfp* reporter with the *wee2* 3’ UTR antisense sequence at the approximate concentration of 200 pg produced strong and consistent GFP signal (Fig. 5 D), similarly to previous studies using GFP reporter assays (Yoshizaki *et al.* 2005). This was in contrast with the results of injections with reporters having *wee2* 3’UTR in the sense orientation, where injections as high as 1600 pg were required for consistent, but faint, GFP signal to be detected (Fig. 5 A). This indicated suppression of *wee2* by endogenous factors binding to its 3’UTR. Observations that embryos with severe deformities would produce intense GFP signal from reduced *gfp-wee2* concentrations provided further evidence that endogenous factors were acting on the *wee2* 3’UTR during zebrafish embryogenesis.

In zebrafish, miRNAs predicted to target the *wee2* 3’ UTR include: miR-30, miR-96, miR-101, miR-137, miR-145, miR-148/152, miR-155/2194, miR-156, as well as miR-25-3p, miR-92a-3p, miR-92b-3p, and miR-363; the four latter ones share the same predicted target site (Agarwal *et al.* 2015). Among all these miRNAs, miR-92a-3p is by far the most abundant in zebrafish unfertilized eggs, while miR-25-3p is also relatively abundant (Presslauer *et al.* 2017), suggesting these miRNAs are likely the endogenous factors acting to suppress *wee2* translation.

### miR-92a-3p suppresses *wee2* translation in early development and promotes cell cycle progress

The present study demonstrates novel information on miRNA functionality in the cell cycle regulation at the onset of embryogenesis. Several miRNAs are known to be involved in cell cycle regulation (Silva Rodrigues *et al.* 2018). In mammalian embryonic stem cells, the conserved miR-17-92 cluster supports cellular reprogramming through inhibiting regulatory elements of the G1 phase and G1/S checkpoint. At the G2/M checkpoint, however, the only miRNA with known functionality is miR-195, which suppresses Wee1 transcripts (Mens and Ghanbari 2018).

Co-injection of the *gfp-wee2* reporter with a target site blocker for the predicted miR-92a-3p binding site at the *wee2* 3’UTR resulted in strong *gfp* expression, even at a low concentration of the *gfp-wee2* reporter (Fig. 6). A significant increase in proportion of GFP-expressing embryos, with significantly increased GFP intensity among those embryos, suggested functionality of that binding site in suppression of *wee2* by endogenous miRNAs. Co-injection of the *gfp-wee2* reporter with a miR-92a-3p mimic resulted in a decrease in the number of GFP-expressing embryos, and a decrease in GFP intensity among those embryos (Fig. 7). Together, it demonstrates that miR-92a-3p is functional in the suppressing maternal *wee2* during the early development of zebrafish.

Previously we observed that the inhibition of maternal miR-92a-3p by injection of a *vivo* MO directly into an ovary induced an irreversible developmental arrest of the resulting embryos at the 1-cell stage (Presslauer *et al.* 2016). In the present study, the delivery of miR-92a-3p MO into 1-cell embryos resulted in a non-lethal phenotype, manifested in a significant proportion of embryos displaying slowed development at 1 dpf. The effect was reversible, and the embryos fully recovered by 2 dpf (Fig. 8). These results demonstrate that the timing of the miR-92a-3p-MO application is decisive for the essentiality of unblocking the suppression of the G2/M checkpoint. Due to lack of miR-92a-3p, a posttranscriptional suppressor, the concentration of Wee2 in miR-92a-3p knockdown oocytes and further zygotes (Presslauer *et al.* 2016) was high enough to block the progression of the first mitotic division; whereas, the miR-92a-3p-MO in the present study was delivered later, into the 1-cell stage embryos. The *wee2* transcript level is rather stable throughout the cleavage stages (Fig. 4), a typical feature for maternal transcripts (Tadros and Lipshitz 2009). In the cell cycle regulation context, it means that the concentration of this transcript per nucleus is reduced roughly by half with every cleavage. Thus, the temporal developmental retardation, observed in the present study, could plausibly be associated with diminishing essentiality of Wee2 in the cell cycle suppression, caused by the decreasing concentration of the Wee2 per blastomere. Taken together, the results of the previous (Presslauer *et al.* 2016) and the current study suggest that miR-92a-3 is essential in promoting the cell cycle progression throughout the first cleavage stages in zebrafish through suppressing translation of CDK1 inhibitors, primarily Wee2.

In conclusion, miR-92a-3p is strongly expressed in both gonadal somatic and germline cells in zebrafish. During the early embryonic development, maternal miR-92a-3p transcripts are co-expressed with their predicted target *wee2*, a tyrosine kinase inhibitor of cell cycle progression. Endogenous miRNAs interact with the *wee2* 3’UTR, and synthetic miR-92a-3p is capable of suppressing *gfp-wee2* translation. Knockdown of miR-92a-3p in zebrafish embryos resulted in delayed development during the first day of embryogenesis, while the developmental arrest at the 1-cell stage was observed in the previous study, when the knockdown was performed in mature ovaries. These results suggest that zebrafish maternal miR-92a-3p has essential role in regulating the cell cycle through suppressing CDK1 inhibitors, primarily Wee2. This role is particularly important for the progression through the first cleavage.

## Methods

### Fish

The zebrafish used in the experiment were from the inbred AB line, which was originally obtained from The Norwegian Zebrafish Platform (zebrafish.no), Norwegian University of Life Sciences (Oslo, Norway), before becoming established at Nord University (Bodø, Norway). The zebrafish were housed in an Aquatic Habitats recirculating system (Pentair, Apopka FL, USA) following standard zebrafish husbandry procedures (Westerfield 2000). Zebrafish nutrition consisted of a daily mix of newly hatched *Artemia* nauplii (Pentair) and SDS zebrafish specific diet (Special Diet Services, Essex, United Kingdom) following the manufacturers recommended feeding regime.

All experimental procedures described in the present study were performed in accordance with the Norwegian Regulation on Animal Experimentation (The Norwegian Animal Protection Act, No. 73 of 20 December 1974) and were approved by the National Animal Research Authority (Utvalg for forsøk med dyr, forsøksdyrutvalget, Norway) General License for Fish Maintenance and Breeding (Godkjenning av avdeling for forsøksdyr) no. 17.

### Sampling

Total RNA was extracted from the ovaries of three mature zebrafish. The fish were first euthanized with 200 mg/L of MS-222 Tricaine; Sigma Aldrich, Oslo, Norway) buffered with equal parts sodium bicarbonate (NaHCO_3_; Sigma Aldrich). After opercular movement had ceased, the fish were decapitated, the gonads were immediately removed, and total RNA was extracted using QIAzol Lysis Reagent (Qiagen, Hilden, Germany) following the manufacturer’s instructions. RNA integrity was assessed using 1% (w/v) agarose gel electrophoresis and was quantified using a NanoDrop ND-1000 (Thermo Fisher Scientific, Saven & Werner AS, Kristiansand, Norway).

Ten-week-old zebrafish were sampled for *in situ* hybridization (ISH). The fish were euthanized as described above; the abdominal cavity was opened, and the specimens were placed directly into pre-chilled Bouin’s solution (4 °C; Sigma Aldrich). The samples were fixated overnight at 4°C before being dehydrated in a gradient series of ethanol washes (from 25 to 100 %) and embedded in paraffin wax. The samples were systematically sectioned (6 μm thickness) using a rotary microtome (Microm HM355S, MICROM International GmbH, Germany) and were mounted on Polysine^®^slides (Thermo Fisher Scientific). Slides were stored at 4 °C for a maximum of one week before use.

Zebrafish embryos were both fixed for whole mount *in situ* hybridization (WISH), and snap-frozen for total RNA extraction. Embryos were produced by allowing single pairs of adult zebrafish to spawn naturally while under observation to ensure accurate knowledge of the time of fertilization. After embryos were collected, they were individually examined using a stereomicroscope and staged according to previously established guidelines (Kimmel *et al.* 1995). For the WISH experiment, zebrafish embryos at the 1-cell, 256-cell, and 25 somite stages were collected and placed directly into pre-chilled 4% w/v paraformaldehyde (PFA; pH 7.0) and fixated overnight at 4 °C. The following day the embryos were repeatedly washed with a chilled solution of 0.1 % Tween-20 in phosphate buffered saline solution (PBST) before WISH. For total RNA extraction, three additional spawning’s were conducted. In each group of embryos, samples were taken at seven developmental stages: activated (5 min post¬fertilization), 1-cell, 4-cell, 32-cell, 256-cell, oblong, and 25 somite. For each sampling, pools of ten embryos were collected and placed directly in QIAzol Lysis Reagent.

### Reverse transcription

Complementary DNA (cDNA) was synthesized from total RNA previously extracted from mature zebrafish ovaries and embryos. Reverse transcription was performed using the Quantitect Reverse Transcription Kit (Qiagen) following the manufacturer’s protocol. For ovaries and for embryos, 1.0 μg and 100 ng total RNA input was used, respectively. Reverse transcription of small RNAs was performed for all embryonic stages using the miRCURY LNA RT kit (Qiagen) using the recommended 10 ng total RNA input.

### *In situ* hybridization of zebrafish gonads

A locked nucleic acid (LNA) oligonucleotide probe antisense for the mature form of miR-92a-3p was designed and produced by Qiagen. The probe sequence was ACAGGCCGGGACAAGTGCAATA, and the probe was double digoxigenin labeled at the 5’ and 3’ ends. As a negative control, a LNA™ Scramble-miR negative control probe was used (GTGTAACACGTCTATACGCCCA).

The ISH procedure was performed in accordance to the miRCURY LNA™ microRNA ISH Optimization Kit (Qiagen) with some modifications. Melting of paraffin was performed for 1 h at 60 °C the day prior to the ISH experiment. Section permeability was improved using 12 μg/mL Proteinase-K (Roche) for 10 minutes at room temperature. LNA probe hybridization was performed with a probe concentration of 40 nM in microRNA ISH buffer (Qiagen). Hybridization occurred at 30 °C below the RNA Tm °C (57 °C for both miR-92a-3p and the scrambled control) for 1 h.

### Whole mount *in situ* hybridization of zebrafish embryos

A cRNA probe for zebrafish *wee1 homolog 2* (*wee2;* accession: NM_001037222) was developed by first amplifying a partial cDNA sequence from zebrafish ovaries which encompassed the majority of the 3’ untranslated region (3’ UTR). Because zebrafish *wee2* has a transcript variant (*wee2-204*; ENSDART00000151974) with an alternative stop codon, we also amplified an additional region for sequencing. PCR conditions and oligonucleotide sequences are summarized in Supplementary Table S1. PCR products of the expected size were excised from a 1 % w/v agarose gel and purified using the QIAquick gel extraction kit according to the manufacturer’s protocol (Qiagen). The cDNA fragment was inserted into the pCR4-TOPO^®^ vector and transformed into OneShot^®^ chemically competent *E. coli* cells. After propagation, the vector was purified using the QIAprep® Miniprep kit (Qiagen) and sequenced in both directions using M13 primers with the Big Dye® Terminator 3.1 (Applied Biosystems) sequencing template preparation method. The sequencing reactions were analyzed at the DNA Sekvenseringslab, University of Tromsø, Norway. The confirmed zebrafish *wee2* cDNA fragment was amplified from the pCR4-TOPO^®^ vector into linear fragments using the M13 primers. No vectors contained the PCR products of the *wee2-204* transcript variant. The extracted PCR product was then used as a template for digoxigenin labeled cRNA probe transcription with T3 (anti-sense) and T7 (sense) RNA polymerases (Roche, Mannheim, Germany). The cRNA probes were purified using LiCl/ethanol precipitation and re-suspended in nuclease-free water before storage at −80 °C.

WISH for miR-92a-3p and *wee2* was performed as previously described with minor adjustments (Hall *et al.* 2003). Embryo permeability was improved using 2.5 μg/mL Proteinase-K (Roche) for 10 min at room temperature. The miR-92a-3p probe hybridization was performed with a probe concentration of 40 nM for 1 h at 57 °C, whereas *wee2* probe hybridization was performed with a concentration of 1.0 μg/μL for 48 h at 70 °C. Bound probe was conjugated to alkaline-phosphatase labelled anti¬DIG antibody (Roche) overnight at a 1:1000 dilution. The colour reactions were performed at room temperature for 1 and 10 h for miR-92a-3p and *wee2*, respectively.

### Quantitative PCR

Transcript quantification was performed using the QX200™ Droplet Digital PCR system (Bio-Rad laboratories, Oslo, Norway) according to the manufacturer’s guidelines. Custom oligonucleotides were designed for *wee2* (Supplementary Table S1), while previously validated oligonucleotides for *ribosomal protein* |13α (rpl13α; accession: NM_001037222) and *actin beta 2* (*actb2*; ENSDART00000055194) were used for comparison (Tang *et al.* 2007b). For small RNAs, miRCURY LNA™ miRNA PCR Assays were designed and synthesized by Qiagen for miR-92a-3p as well as miR-19d-3p (miRBase accession: MIMAT0001785), and miR-192-5p (miRBase accession: MIMAT0001275); the latter two were used for comparison, as they had previously been detected in zebrafish eggs (Presslauer *et al.* 2017). For both mRNA and small RNA, 20x diluted cDNA inputs were used.

### miR-92a-3p target validation

Zebrafish miR-92a-3p target validation was performed through a series of microinjection experiments. First, the fragment of *wee2* 3’UTR, previously amplified for WISH, was excised from the pCR4-TOPO^®^ vector using EcoRI digestion (Thermo Fisher Scientific). After gel purification, the fragment was ligated into a pGEM-T-Easy vector (Promega) containing the *hrgfp* coding region (Yoshizaki *et al.* 2005). To ensure the proper insert orientation, several colonies were propagated and sequenced as previously described. Plasmids containing both the sense and antisense *wee2* 3’UTR were linearized using NdeI (Thermo Fisher Scientific) and *in vitro* transcription was performed using the Message Machine T7 Kit (Thermo Fisher Scientific). The transcribed RNAs were purified using LiCl precipitation and re-suspended in 20 μL nuclease-free water before being aliquoted and stored at −80 °C. The calculated stock concentrations of *gfp-wee2* and *gfp-wee2-antisense* were 3200 ng/μL and 1000 ng/μL, respectively (NanoDrop ND-1000).

In order to determine the optimal concentration of chimeric RNA for injection, to obtain strong GFP signal, the *gfp-wee2* and *gfp-wee2-antisense* chimeric mRNAs were diluted in freshly made and chilled embryo medium (Westerfield 2000) before injecting into naturally spawned 1-cell zebrafish embryos approximately 30 to 60 min post fertilization. *gfp-wee2* injection concentrations were 3200, 1600, 800, and 200 ng/μL compared to 1000 and 200 ng/μL for the *gfp-wee2-antisense* control. For all injections, 1 nL volume was injected into the yolk directly below the blastodisc using an IM-300 microinjector (Narishige, London, UK), resulting in respective dosages of 200 pg to 3200 pg mRNA. After injection, embryos were transferred to a fresh solution of embryo medium for incubation at 28.5 °C. All observations were performed using an AxioZoom V.16 microscope (Carl Zeiss, Göttingen, Germany) and Zen Pro (2012; Carl Zeiss) imaging software.

The miR-92a-3p target site blocker assays were performed using a miRCURY LNA Power Target Site Blocker (TSB) specific for the miR-92a-3p binding site within the zebrafish *wee2* 3’UTR. The TSB (5’-AT<u>TATTGCAC</u>CCAGTGCC-3’: the miR-92a-3p target site is underlined) was designed and produced by Qiagen, with an additional scrambled TSB to act as a negative control (5’-TAACACGTGTATACGCCCA-3’). Upon receipt, the lyophilized TSBs were re-suspended in nuclease free water at a stock concentration of 50 μM and aliquoted for further use. The TSB assays were optimized for a 1 nL injection volume consisting of 320 ng/μL *gfp-wee2* and 1.0 μM TSB in freshly prepared embryo medium. Microinjections and general observations were made as previously described. The GFP mean signal intensity was measured using the ZEN Measurement Module (Zeiss). For each measurement, contours were traced to encompass the head, brain, and eye region of the embryo; these regions were the most consistent in their GFP expression.

*mirVana*^®^ miRNA mimic for zebrafish miR-92a-3p was designed and produced by Thermo Fisher Scientific. The lyophilized mimic was re-suspended in nuclease-free water to a stock concentration of 100 μM and aliquoted for further use. The effect of miR-92a-3p mimic on *gfp-wee2* expression was observed by a series of 1.0 nL injections consisting of 5.0 μM mimic and 800, 1600, or 3200 ng/μL *gfp-wee2*. Observations and measurements were made as previously described.

### miR-92a-3p knockdown

miR-92a-3p knockdown was performed using a vivo-morpholino (VMO) complementary to the dre-miR-92a-3p guide strand (5’-TACAGGCCGGGACAAGTGCAATACC-3’) designed and produced by Genetools LLC (Philomath, OR, USA) with an additional 5-base mismatch VMO (5’-TA<u>G</u>A<u>C</u>GCCG<u>C</u>GACAA<u>C</u>T<u>C</u>CAATACC-3’; underlined letters indicate the mismatches) as a negative control. Microinjections and embryo staging were performed as previously described. Treatments consisted of 5.0 μM VMO and were rescued with an equal concentration of miR-92a-3p mimic. Embryos were examined for phenotypes at 1 and 2 days post-fertilization.

### miR-92a-3p target prediction

Targets were predicted for zebrafish miR-92a-3p (miRBase accession: MIMAT0001808) using both TargetScanFish (release 6.2; http://www.targetscan.org/fish_62/) and miRmap (http://mirmap.ezlab.org/) (Vejnar and Zdobnov 2012; Agarwal *et al.* 2015). Potential targets were filtered using the Kyoto Encyclopedia of Genes and Genomes (KEGG; http://www.genome.jp/kegg/pathway.html) pathway database to select those involved in the zebrafish cell cycle pathway (Kanehisa and Goto 2000).

### Statistical analyses

Expression values from ddPCR analyses were log transformed and the effect of developmental advancement on transcript abundance was estimated using ANOVA. The Brown-Forsythe test was used to evaluate homogeneity of variances. Tukey’s HSD post-hoc test was used to estimate differences in transcript abundance among developmental stages. For GFP-reporter assays, the mean intensity values were tested with F-test for two equal variances before evaluating the effect of the treatment using the two-tailed Student’s *t-*test. Percentage data (frequency of GFP expressing embryos and frequency of embryos with delayed development) were arcsin square root transformed. ANOVA was used to analyse differences in development between miR-92a-3p knock-down fish, the mismatch control, miR-92a-3p mimic, and the rescue treatment (morpholino and mimic combination). Multiple comparison analysis was performed using the Tukey-Kramer test. All effects were considered significant at a *p*-value of 0.05 of less (Zar 1999).

## Acknowledgements

We thank the Mørkvedbukta Research Station technicians, in particular Cesilie Amundsen and Katrine Klippenberg, for their assistance in zebrafish husbandry, and Heidi Ludviksen for aid in laboratory operations.

## Supplementary Files

**Supplementary Table S1. PCR reaction conditions and primer sequences used in the study.**

**Supplementary File S1. Predicted cell cycle targets for miR-92a-3p (miRBase accession: MIMAT0001808) using both TargetScanFish and miRmap, and compared to the zebrafish cell cycle KEGG pathway.**

**Supplementary File S2. Quantitative PCR of zebrafish maternal miRNAs and mRNAs during early embryogenesis.**

